# AptCompare: optimized de novo motif discovery of RNA aptamers via HTS-SELEX

**DOI:** 10.1101/413757

**Authors:** Kevin R. Shieh, Christina Kratschmer, Keith E. Maier, John M. Greally, Matthew Levy, Aaron Golden

## Abstract

**Summary:** High-Throughput Sequencing can enhance the analysis of aptamer libraries generated by the Systematic Evolution of Ligands by EXponential enrichment (HTS-SELEX). Robust analysis of the resulting sequenced rounds is best implemented by determining a ranked consensus of reads following the processing by multiple aptamer detection algorithms. Whilst several such approaches have been developed to this end, their installation and implementation is problematic. We developed AptCompare, a cross-platform program that combines six of the most widely used analytical approaches for the identification of RNA aptamer motifs and uses a simple weighted ranking to order the candidate aptamers, all driven within the same GUI-enabled environment. We demonstrate AptCompare’s performance by identifying the top-ranked candidate aptamers from a previously published selection experiment in our laboratory, with follow-up bench assays demonstrating good correspondence between the sequences’ rankings and their binding affinities.

**Availability and Implementation:** The source code and pre-built virtual machine images are freely available at https://bitbucket.org/shiehk/aptcompare.

## INTRODUCTION

Aptamers are oligonucleotides which possess specific binding affinity for their intended targets. To date, aptamers with high affinity and selectivity have been reported for various targets, including small molecules, proteins, and cell-surface receptors (Stoltenburg et al., 2007). Compared to antibodies, aptamers have several advantages: they can be synthesized in vitro, are inexpensive to produce, and are not immunogenic (Yan and Levy, 2009).

The technique by which aptamers are generated is known as in vitro selection or SELEX (System Evolution of Ligands by EXponential enrichment) (Ellington and Szostak, 1990; Tuerk and Gold, 1990). In this approach, a randomized sequence pool is subjected to multiple iterations of binding to a molecular target, elution from that target, and amplification. Between five and fifteen rounds of SELEX are typically required to yield detectable function which can be attributed to individual clones following sequence analysis and subsequent assays.

Traditionally accomplished by molecular cloning, advances in DNA sequencing technology have increased the sampling resolution of the RNA species and enabled a quantitative analysis of the final, enriched population in each selection round. As aptamers bind to their targets, individual molecules or families of similar sequences harboring one or more ‘active’ motifs are expected to become enriched throughout the course of selection. When analyzed by high-throughput sequencing (HTS), reads from each round of the selection can be used to identify the overrepresented subsets, and the resulting consensus aptamer sequences can be determined. These can then be modified or minimized at the bench for improved efficiency in synthesis and binding.

To date several programs have been developed to assist with motif discovery from aptamer selections following HTS-SELEX, each with different modes of implementation, installation, and overall performance. In this article, we present a convenient means for researchers to conduct a comprehensive comparison of six of the most widely used methods of analysis using a software environment called AptCompare. AptCompare automates the preprocessing and analysis of HTS-SELEX sequencing data and uses a simple scheme to rank the weighted scores from all six methods per processed round within the same GUI environment.

## IMPLEMENTATION

AptCompare combines several approaches for de novo motif discovery in aptamer selections. The first approach is a frequency or sequence counting script written in Perl, which performs a simple ranking of the most frequent sequences in an aptamer pool that is often the first step in any such analysis. The other five programs programs are: AptaCluster (AptaTools) (Hoinka et al., 2014), FASTAptamer (Alam et al., 2015), MPBind (Jiang et al., 2014), APTANI (Caroli et al., 2016), and RNAmotifAnalysis (Ditzler et al., 2013). Of these six approaches, the first four are sequence-based, whereas the last two incorporate RNA folding programs to predict secondary structure. All of these approaches, their features, and their requirements are summarized in Supplementary Table S1.

Designed to be cross-platform, AptCompare was developed primarily in Python, with the GUI written using the PyForms and PyQt4 libraries. The individual programs, however, may have software dependencies, including Unix utilities, MySQL, and third-party components. Therefore, we have also provided a virtual machine image and an Amazon Machine Image (AMI) to facilitate ‘turn-key’ configuration and deployment on new systems.

In order to make equivalent comparisons, we have designed AptCompare to require minimal user input and configuration in most situations. This ‘basic’ mode executes the analysis with default parameters. Advanced users can configure specific parameters and run the individual programs. Following the completion of the analysis, the results can be exported into a tab-delimited format and saved as a spreadsheet. The results are sorted by the weighted ranking, the mean rank calculated by all six methods. AptCompare also sums the number of programs (out of six) that select a given motif, corresponding to a ‘measure of belief’ in that motif model.

A preprocessing step converts raw FASTQ, FASTA, or sequence files to all necessary formats for each of the individual programs. It also relies on cutadapt (Martin, 2011) to remove the flanking constant regions from the sequences. Because we are primarily interested in aptamer motifs, AptCompare uses an initial hierarchical clustering step in the FASTAptamer package to identify cluster seeds, the representative sequences of each cluster. The remainder of the analysis is conducted using these cluster seeds, significantly enhancing overall performance.

## RESULTS

We evaluated the performance of AptCompare using the data set from an aptamer selection against the human transferrin receptor (hTfR) previously conducted in our laboratory (Wilner et al., 2012; Maier et al., 2016). The original selection, which did not use HTS, identified three aptamers, all of which share a core motif, characterized by two asymmetric internal loops and a longer motif in one of the loops (Supplementary Figure S1(a)). We sequenced each round of this selection using HTS and ran AptCompare on the resulting reads from each round. Twenty-seven unique ranked candidate aptamers were identified in total (details in Supplementary Data). Multiple sequence alignment reveals that nearly all share the common nonamer GATCA[AT]TNC, which was the motif identified in the previous selection (Supplementary Figure S1(b)).

We synthesized all twenty-seven candidate aptamers and measured their binding affinity via flow cytometry binding assays (see the Supplementary Data for methods). The binding affinities were measured by estimating the apparent dissociation constant, K_d_. Our results demonstrate that the binding affinity correlates well with the rankings of the aptamers (Supplementary Tables S2 and S3) generated using all five programs, with no one program completely outperforming any of the others. Interestingly, even the simplest method, sequence counting, proved highly effective. Thus, programs employing structural prediction or statistical modeling that are generally significantly more time-consuming to implement may only provide a limited amount of additional information.

We have designed AptCompare with experimentalists in mind, who need to analyze the results from aptamer selections derived from HTS-SELEX and wish to base their validation assays on a comprehensive study of the derived read libraries. AptCompare conveniently performs preprocessing, clusters sequences, and compares the results from six of the most widely used approaches encompassing a variety of sequence- and secondary structure-based techniques. It calculates a weighted ranking and the number of appearances, which are analogous to the significance of a motif and a measure of belief in that motif, respectively. Validation assays on novel candidate aptamers generated from a previously published study demonstrated a clear correlation between the measured binding affinities and their rankings. AptCompare was designed to be a modular program built on Python that integrates various data analysis methods used for RNA aptamers. It is a model for a meta-comparison of analytical methods, especially if no de facto standard exists in the field, and combined with its ‘turn key’ design, makes it a highly effective tool for RNA aptamer discovery.

## ACKNOWLEDGMENTS

The authors acknowledge the support and advice of Myles Akabas, MD, PhD.

